## Supplementary tables and figures for "How do gepotidacin and zoliflodacin stabilize DNA-cleavage complexes with bacterial type IIA topoisomerases? 1. Experimental definition of metal binding sites"

Table S1: Re-refinement statistics on 2xcs (2.1Å), 5cdm (2.5Å), 5cdr (2.65Å) and 5iwi (1.98Å).

Table S2: Data collection statistics - 9fz6 (2.58Å - anomalous data set).

Table S3: Refinement statistics - 9fz6 (2.58Å - anomalous data set).

Supplementary reference

Figure S1: A simplified schematic illustrating a mechanism of action of DNA gyrase.

Figure S2: A comparison of superposed and non-superposed structures.

Figure S3: The active site $Mn^{2+}$ coordination geometry in the 2.1Å GSK299423 structure (pdb code: 2xcs).

Figure S4: Active site $Mn^{2+}$ coordination geometry in 2.5Å QPT-1 structure (pdb code: 5cdm) compared with a 2.16Å human top IIb structure with etoposide (3qx3).

Figure S5: The originally deposited coordinates of 2xcs contained an error at a crystal contact near the Xtal contact $Mn^{2+}$ ion.

Table S1. Re-refinement statistics on 2xcs (2.1Å), 5cdm (2.5Å), 5cdr (2.65Å) and 5iwi (1.98Å)

| | **1.**<br>**3' (A) site $Mn^{2+}$**<br>Xtal contact $Mn^{2+}$<br>-<br>**GSK299423** | **5.**<br>**Y (B) site metal**<br>Xtal contact $Mn^{2+}$<br><br>**QPT-1** | **7.**<br>**Y(B) site metal**<br>Xtal contact $Mn^{2+}$<br>E1999, F1999 $Mn^{2+}$<br>No inhibitor | **8.**<br>**3'(A)+Y(B)-sites**<br>**(occupancies two 0.57 and two 0.43)**<br>**GSK945237** |
|---|---|---|---|---|
| **New PDB code** | 2xcs-v2 | 5cdm-v2 | 5cdr-v2 | 5iwi-v2 |
| **Resolution range (Å)** | 24.0-2.1 (2.154-2.10) | 18.0-2.5 (2.564-2.50) | 18.0-2.65 (2.717-2.65) | 18.0-1.98 (2.031-1.980) |
| **Completeness (%)** | 98.53 (98.04) | 98.39 (98.80) | 99.02 (99.91) | 99.17 (97.60) |
| **No. of reflections, work set** | **113170 (8275)** | 65867 (4826) | 55539 (4020) | 134611 (9674) |
| **No. of reflections, test set** | 2907 (171) | 3508 (265) | 2372 (190) | 2759 (199) |
| **Final Rcryst** | 0.167 (0.247) | 0.175 (0.166) | 0.163 (0.232) | 0.157 (0.271) |
| [Previous Rcryst ] | [0.184 (0.235)] | 0.165 | [0.189. (0.233)] | [0.166 (0.277)] |
| **Final Rfree** | 0.199 (0.285) | 0.196 (0.212) | 0.208 (0.266) | 0.189 (0.281) |
| [Previous Rfree ] | [0.214 (0.277)] | 0.192 | [0.210 (0.254)] | [0.202 (0.295)] |
| **Refined twin fractions** | Not twinned | 0.215/0.785 | Not twinned | Not twinned |
| **Estimated coord. error (Å)**** | 0.16/0.14/0.10 | 0.08/0.04/0.12 | 0.69/0.27/0.20 | 0.136/0.124/0.097 |
| [Previous Est. coord err.] | [0.18/0.15/0.11] | [Phenix refined] | [0.61/0.255/?] | [0.141/0.131/0.106] |
| **No. of non-H atoms (total)** | 11791 | 11619 | 11740 (11578*) | 11981 (10913*) |
| **Protein** | 10175 | 10405 | 10258 (10230*) | 10011 (9758*) |
| **DNA** | 775 | 769 | 739 | 797 |
| **GSK299423*/QPT-1 /None/GSK945237** | 33(66)* | 54 | None | 33 (66)* |
| **$Mn^{2+}$ ions full/partial***** | 2 /1 | 2/1(0.4) | 2/3(0.4) | 4*** /1(0.3) |
| **Waters** | 826 | 344 | 703 | 1115 (1099*) |
| **Partial BTB with Mn** | 14 (0.65) | 14 (0.4) | 14 (0.3) | 14 (0.3) |
| **RMS deviations (Refmac)** | | | | |
| **Bonds (Å)** | 0.006 | 0.007 | 0.006 | 0.007 |
| **Angles (°)** | 1.585 | 1.457 | 1.561 | 1.457 |
| **Average B factors (Å²)** | | | | |
| **Protein** | 29.5 | 52.0 | 49.1 | 33.8 (33.7) |
| **DNA** | 28.2 | 45.0 | 53.8 | 31.5 |
| **GSK299423*/QPT-1/ None/GSK945237*** | 30.8 (66*) | 37.6 | - | 30.0 (66*) |
| **$Mn^{2+}$ ions full/partial***** | 21.9 / 23.3 | 36.7/56.5 | 37.4/56.3 | 25.0 /35.8 |
| **Waters** | 32.6 | 44.2 | 42.0 | 43.2 (43.3) |
| **Partial BTB with Mn** | 30.3 | 55.4 | 37.2 | 40.5 |
| **Ramachandran plot** | | | | |
| **Favored regions** | 97.45 | 97.3 | 97.7 | 97.6 |
| **Additionally allowed** | 2.52 | 2.55 | 2..30 | 2.33 |
| **Outliers** | 0.07 | 0.15 | 0.0 | 0.07 |

*GSK299423 and GSK945237 sit on twofold axis of complex. Both compounda contain 33 atoms but these are modelled in two equivalent positions. In the 1.98Å structure with GSK945237 the DNA is disorderd around the twofold axis. Setting occupancy 0.57 = 1.00 gives 11981 atoms with CCP4 program 'bfactor' (the number in brackets is without resetting occupancy from 0.57 to 1.00)

** Estimated coord error = Refmac output = Estimated standard uncertainty based on R VALUE, FREE R VALUE, and MAXIMUM LIKELIHOOD (note both R and freeR values are lower for twinned data - so Maximum likelihood value is used in Supplementary Figures 1 and 2).

*** The metal ions at the two active sites in each complex have full occupancy; metals at other sites have lower occupancies. In the GSK945237 complex electron density is observed for both positions at each active site due to static disorder.

Table S2. Data collection statistics - 9fz6 (2.58Å - anomalous data set).

| | Anom-2.58 |
|---|---|
| **PDB code** | 9FZ6 |
| **Diffraction source** | I23, DLS |
| **Wavelength (Å)** | 1.33 |
| **Resolution range (Å)** | 57.61-2.58 (2.62-2.58) * |
| **Space group** | $P6_1$ |
| **Unit cell** | 93.63, 93.63, 410.87, 90, 90, 120 |
| **Total reflections** | 3,520,698 (59,593) * |
| **Unique reflections** | 63,516 (3132) * |
| **Multiplicity** | 55.4 (19.0) * |
| **Completeness (%)** | 100.0 (100.0) * |
| **Mean I/sigma(I)** | 22.2 (1.1) * |
| **Wilson $B_{factor}$** | 60.3 |
| **$R_{merge}$** | 0.231 (2.924) * |
| **$R_{meas}$** | 0.233 (2.805) * |
| **$R_{pim}$** | 0.031 (0.687) * |
| **$CC_{1/2}$** | 1.000 (0.411) * |

* Numbers in brackets are in the outer (2.62-2.58 Å) resolution shell.

Table S3. Refinement statistics - 9fz6 (2.58Å - anomalous data set).

| | **Anom-2.58** |
|---|---|
| **PDB code** | 9FZ6 |
| **Resolution range (Å)** | 57.72-2.58 (2.62-2.58) |
| **Completeness (%)** | 99.68 (96.37) |
| **No. of reflections, working set** | 63403 (2658) |
| **No. of reflections, test set** | 3158 (127) |
| **Final $R_{cryst}$** | 0.1401 (0.218) |
| **Final $R_{free}$** | 0.1997 (0.346) |
| **Cruickshank DPI (Å)*** | |
| **No. of non-H atoms (total)** | 11966 |
| **Protein** | 11553 |
| **DNA** | 801 |
| **Other ligands (Mn, glycerol etc.)** | 75 |
| **Waters** | 338 |
| **RMS deviations** | |
| **Bonds (Å)** | 0.008 |
| **Angles (°)** | 1.28 |
| **Average B factors (Å$^2$)** | |
| **Protein** | 66.5 |
| **DNA** | 71.5 |
| **Mn (5)*** | 71.7 |
| **Catalytic Mns (2 - full occupancy)** | 55.1 |
| **Other Mns (3 - occcupancy 0.2, 0.6, 0.5)** | 94.0 |
| **Other ligands (glycerol, BisTris etc)** | 103.9 |
| **Waters **** | 58.6 |
| **Ramachandran plot** | |
| **Favored regions** | 94.75% |
| **Additionally allowed** | 5.10% |
| **Outliers** | 0.15% |

*** The Cruickshank DPI (Å) was calculated by the Online_DPI server [1].**

*** The two catalytic metals had 100% occupancy (Y site), the other three Mn ions had partial occupancy.**

**** Waters were placed where there are waters in higher resolution structures (e.g. 5CDM and 5IWI).**

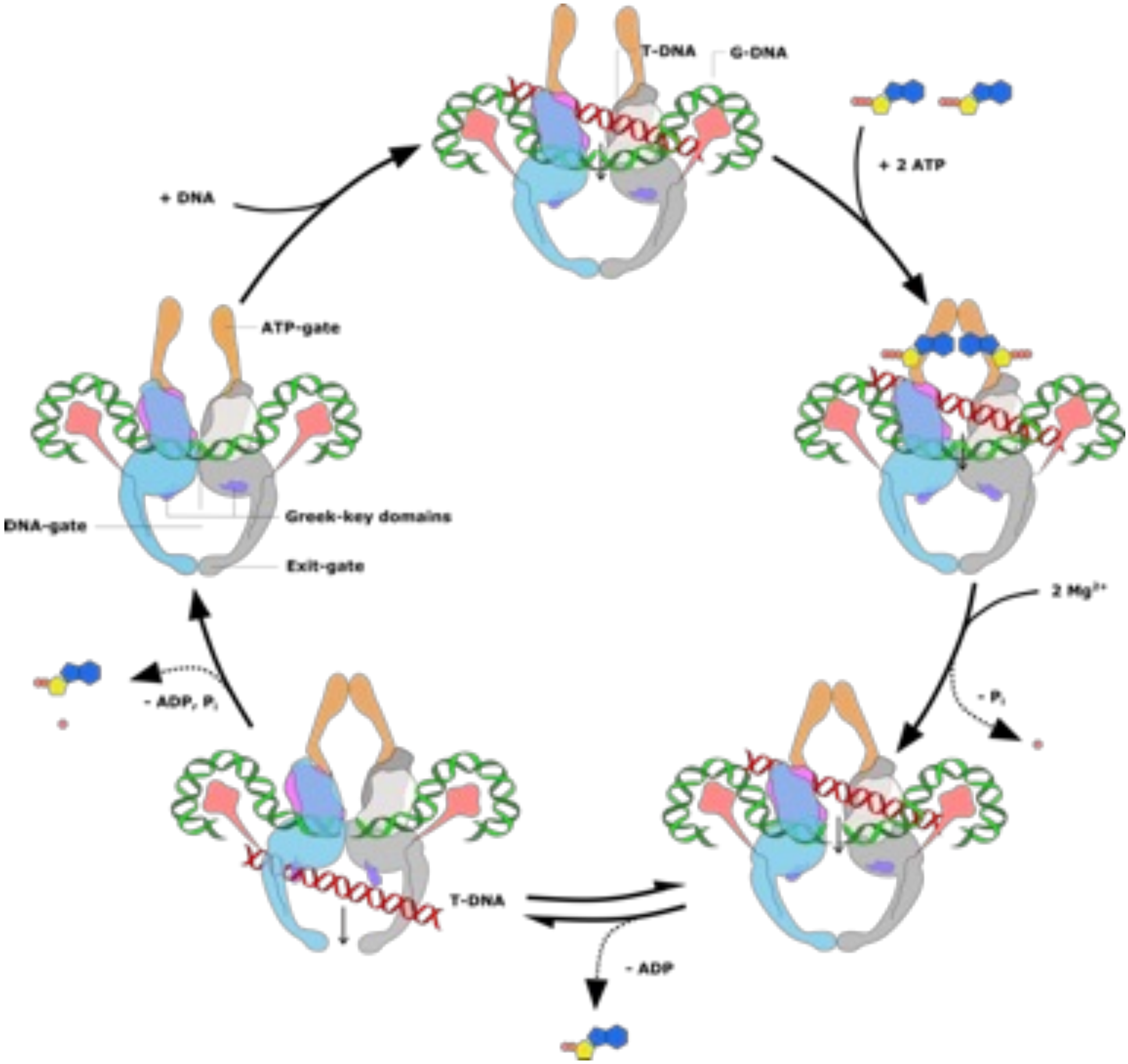


**Supplementary Figure 1: A simplified schematic illustrating a mechanism of action of DNA gyrase.** ATP binding to ATPase domain result can cause in dimerization and T-DNA (red) may be trapped. The G-DNA (green) is transiently cleaved by the two active site tyrosine's from GyrA (WHD). This cleavage allows for the passage of the T-DNA through the G-DNA and enzyme. In this scheme the Greek key domains are not involved in controlling cleavage of the G-DNA segment. Note DNA gyrase has been shown to be capable of catalyzing at least four reactions: (i) the ATP-dependent introduction of negative supercoils into DNA, (ii) the ATP-dependent relaxation of positively supercoiled DNA (iii) the ATP-dependent relaxation of negatively supercoiled DNA and (iv) the ATP-independent relaxation of negatively supercoiled DNA.

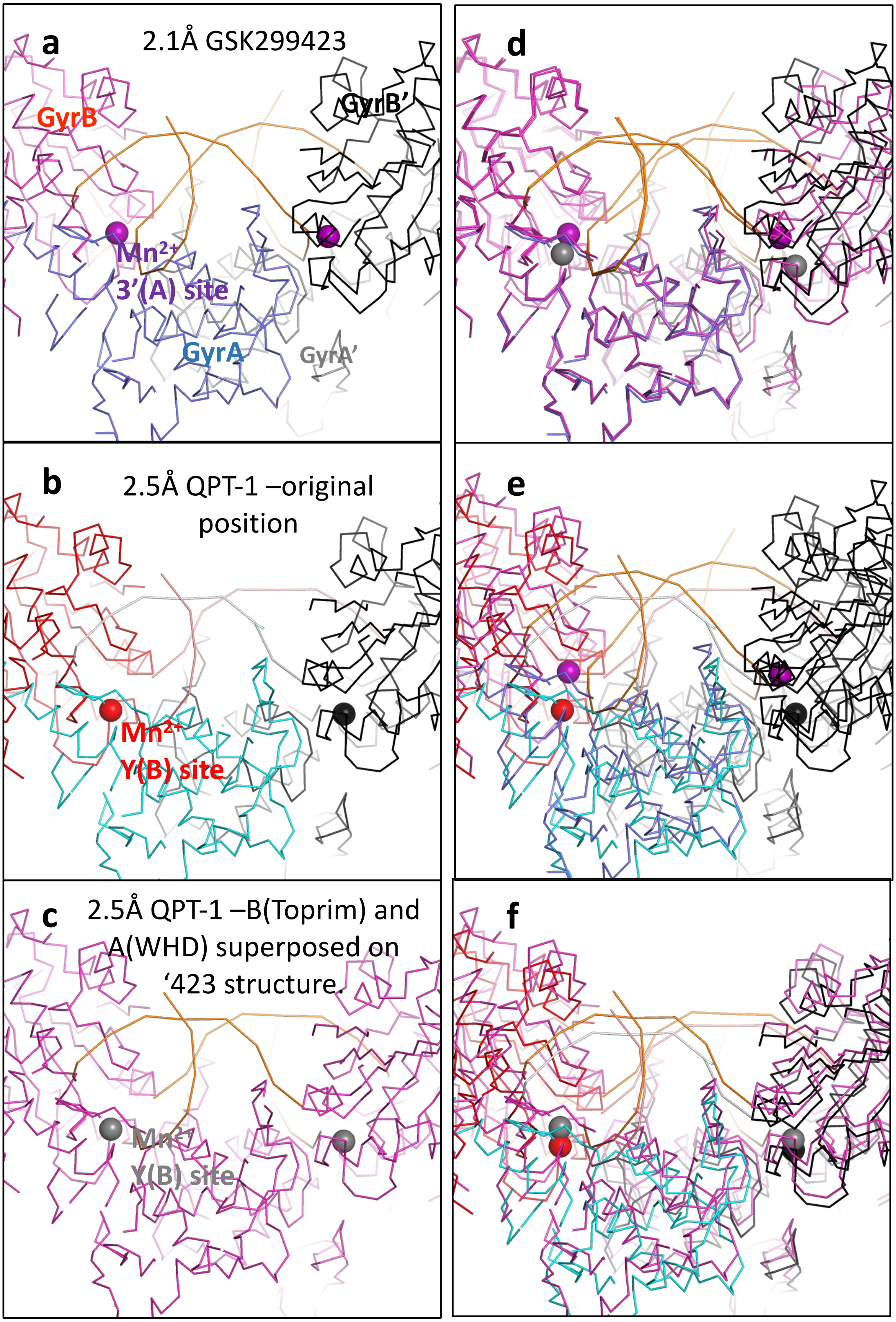


**Supplementary Figure 2: A comparison of superposed and non-superposed structures.**

**(a)** A section through the 2.1Å GSK299423 structure – in ribbon representation. The catalytic manganese ions at the 3'(A) sites are shown as spheres**. (b and c)** Similar sections through the 2.5Å QPT-1 structure – in ribbon representation. The catalytic manganese ions at the Y(B) sites are shown as spheres. (**d**) structures from a and c are shown, note the superposition is OK for the GyrB and GyrA subunits but it is not good for the GyrB' and GyrA' subunits (**e**) structures from a and b are shown (**f**) structures from b and c are shown. The structure superposition shown in c and d was carried out with the program Lsqkab from CCP4 as described in Chan *et al.*, 2015 'Structural basis of DNA gyrase inhibition by antibacterial QPT-1, anticancer drug etoposide and moxifloxacin. *Nat. Commun* **2015**, *6*, 10048, doi:ncomms10048 [pii];10.1038/ncomms10048 [doi].

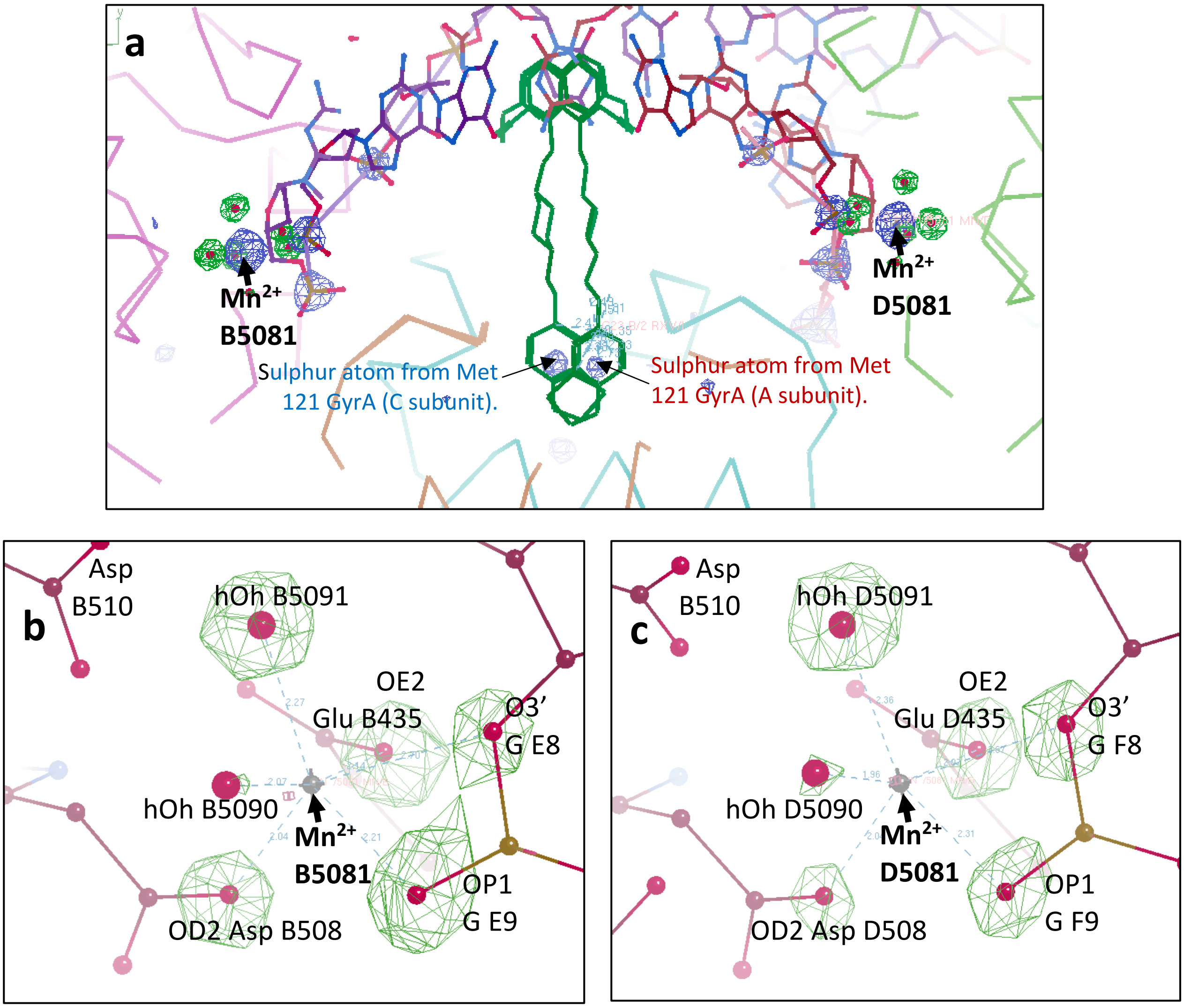


**9.5 sigma omit map – showing 3' (A) site coordination geometry for $Mn^{2+}$ B5081.**

| Atom 1 | Atom 2 | Distance (est. err) |
|---|---|---|
| $Mn^{2+}$ B5081. | OD2 Asp B508 | 2.04Å (0.10Å) |
| $Mn^{2+}$ B5081. | hOh B5090 | 2.07Å (0.10Å) |
| $Mn^{2+}$ B5081. | hOh B5091 | 2.27Å (0.10Å) |
| $Mn^{2+}$ B5081. | OE2 Glu B435 | 2.14Å (0.10Å) |
| $Mn^{2+}$ B5081. | O3' Guanine E8 | 2.70Å (0.10Å) |
| $Mn^{2+}$ B5081. | OP1 Guanine E9 | 2.21Å (0.10Å) |

| Atom 1. | Diff. map peak | Distance (error) |
|---|---|---|
| $Mn^{2+}$ B5081. | on OD2 Asp B508 | 2.07Å (0.14Å) |
| $Mn^{2+}$ B5081. | on hOh B5090 | 2.00Å (0.10Å) |
| $Mn^{2+}$ B5081. | on hOh B5091 | 2.26Å (0.04Å) |
| $Mn^{2+}$ B5081. | on OE2 Glu B435 | 2.15Å (0.14Å) |
| $Mn^{2+}$ B5081. | on O3' Guanine E8 | 2.80Å (0.14Å) |
| $Mn^{2+}$ B5081. | on OP1 Guanine E9 | 2.25Å (0.09Å) |

**9.5 sigma omit map – showing 3' (A) site coordination geometry for $Mn^{2+}$ D5081.**

| Atom 1 | Atom 2 | Distance (est. err) |
|---|---|---|
| $Mn^{2+}$ D5081. | OD2 Asp D508 | 2.04Å (0.10Å) |
| $Mn^{2+}$ D5081. | hOh D5090 | 1.96Å (0.10Å) |
| $Mn^{2+}$ D5081. | hOh D5091 | 2.36Å (0.10Å) |
| $Mn^{2+}$ D5081. | OE2 Glu D435 | 2.07Å (0.10Å) |
| $Mn^{2+}$ D5081. | O3' Guanine F8 | 2.67Å (0.10Å) |
| $Mn^{2+}$ D5081. | OP1 Guanine F9 | 2.31Å (0.10Å) |

| Atom 1. | Diff. map peak | Distance (error) |
|---|---|---|
| $Mn^{2+}$ D5081. | on OD2 Asp D508 | 2.06Å (0.06Å) |
| $Mn^{2+}$ D5081. | on hOh D5090 | 1.87Å (0.09Å) |
| $Mn^{2+}$ D5081. | on hOh D5091 | 2.38Å (0.11Å) |
| $Mn^{2+}$ D5081. | on OE2 Glu D435 | 2.06Å (0.08Å) |
| $Mn^{2+}$ D5081. | on O3' Guanine F8 | 2.73Å (0.06Å) |
| $Mn^{2+}$ D5081. | on OP1 Guanine F9 | 2.37Å (0.13Å) |

**Supplementary Figure 3. The active site $Mn^{2+}$ coordination geometry in the 2.1Å GSK299423 structure (pdb code: 2xcs).** **(a)** A view of the two active sites in the structure. A final omit map in which oxygens coordinating metal ions have been omittted (see panels b and c). Shown is the 2fo-fc 6 sigma (blue) and the fo-fc 9.5 sigma (green). (**b**) The $Mn^{2+}$ ion B5081, showing OMIT map peaks – 9.5 sigma green. Underneath the panel are shown the distances in the refined coordinates. The top six distances are from refined atoms in coordinates (est. err = ESU BASED ON MAXIMUM LIKELIHOOD from Refmac). The bottom six distances are from the refined metal atom to equivalent peaks in the difference map – here the error is estimated as the difference in distance from refined atom to the peak in the difference map (waters in new molecule in difference map and positions refined in coot (Emsley, P.; Lohkamp, B.; Scott, W.G.; Cowtan, K. Features and development of Coot. *Acta Crystallogr. D. Biol. Crystallogr* **2010**, *66*, 486-501, doi:S0907444910007493 [pii];10.1107/S0907444910007493 [doi] ). (**c**) The $Mn^{2+}$ ion D5081, showing OMIT map peaks – 9.5 sigma green. Distances below as in panel b.

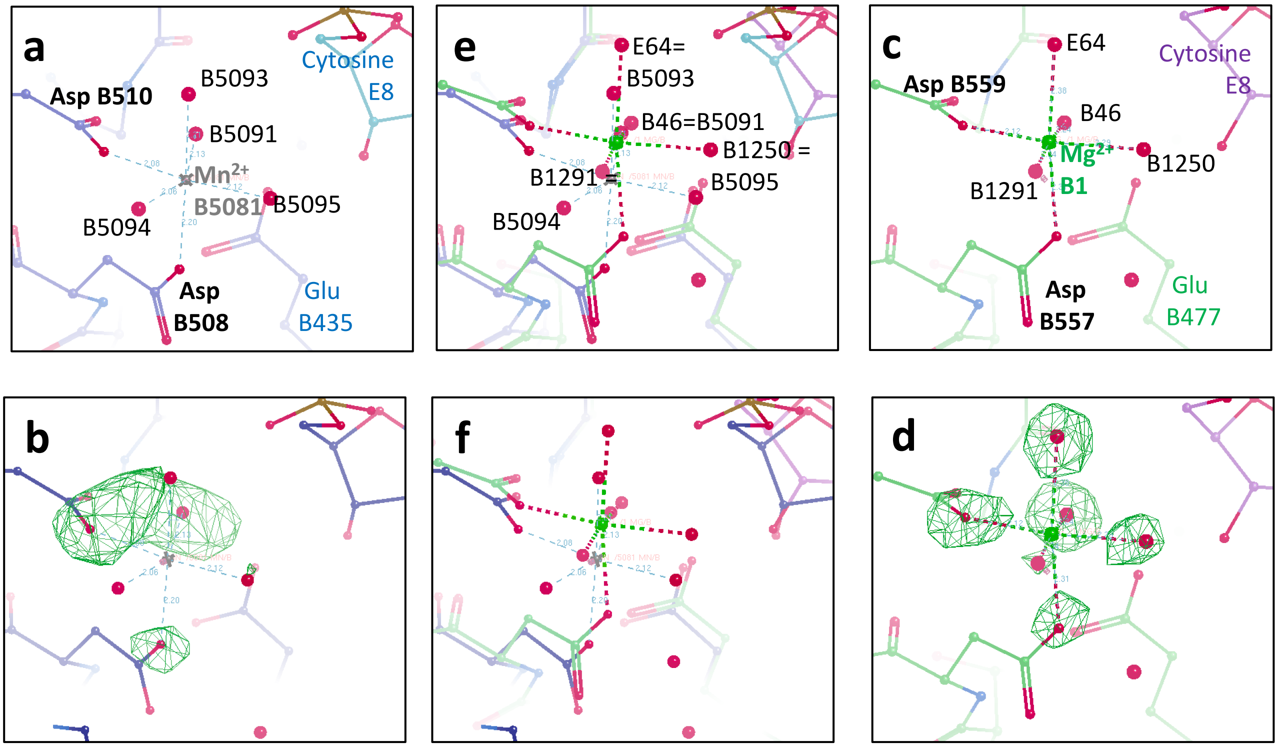


**5 sigma omit map – from 2.5Å 5cdm omit map**

| Atom 1 | Atom 2 | Distance (est. err) |
|---|---|---|
| $Mn^{2+}$ B5081. | OD2 Asp B508 | 2.20Å (0.12Å) |
| $Mn^{2+}$ B5081. | OD2 Asp B510 | 2.08Å (0.12Å) |
| $Mn^{2+}$ B5081. | hOh B5091 | 2.13Å (0.12Å) |
| $Mn^{2+}$ B5081. | hOh B5093 | 2.21Å (0.12Å) |
| $Mn^{2+}$ B5081. | hOh B5094 | 2.06Å (0.12Å) |
| $Mn^{2+}$ B5081. | hOh B5095 | 2.12Å (0.12Å) |

| Atom 1. | Diff. map peak (height) | Distance (error) |
|---|---|---|
| $Mn^{2+}$ B5081. | on OD2 Asp B508 (5.9σ) | 2.36Å (0.19Å) |
| $Mn^{2+}$ B5081. | on OD2 Asp B510 (7.5σ) | 2.11Å (0.55Å) |
| $Mn^{2+}$ B5081. | on hOh B5091 (7.7σ) | 2.20Å (0.29Å) |
| $Mn^{2+}$ B5081. | on hOh B5093 (abs.) | 2.21Å (0.12Å) |
| $Mn^{2+}$ B5081. | on hOh B5094 (abs.) | 1.66Å (0.63Å) |
| $Mn^{2+}$ B5081. | on hOh B5095 (5.1σ) | 2.11Å (0.37Å) |

**7σ omit map – showing Y(B) site coordination geometry for $Mg^{2+}$ in 2.16Å human etoposide structure (3qx3)**

| Atom1 | Atom 2 | Distance (est. err) |
|---|---|---|
| $Mg^{2+}$ B1. | OD2 Asp B557 | 2.31Å (0.25Å) |
| $Mg^{2+}$ B1. | OD2 Asp B559 | 2.12Å (0.25Å) |
| $Mg^{2+}$ B1. | hOh B46 | 2.24Å (0.25Å) |
| $Mg^{2+}$ B1. | hOh E64 | 2.38Å (0.25Å) |
| $Mg^{2+}$ B1. | hOh B1291 | 2.34Å (0.25Å) |
| $Mg^{2+}$ B1. | hOh B1250 | 2.29Å (0.25Å) |

| Atom 1. | Diff. map peak | Distance (error) |
|---|---|---|
| $Mg^{2+}$ B1. | on OD2 Asp B557 | 2.18Å (0.18Å) |
| $Mg^{2+}$ B1. | on OD2 Asp B559 | 2.08Å (0.07Å) |
| $Mg^{2+}$ B1. | on hOh B46 | 2.15Å (0.10Å) |
| $Mg^{2+}$ B1. | on hOh E64 | 2.30Å (0.15Å) |
| $Mg^{2+}$ B1. | on hOh B1291 | 2.09Å (0.26Å) |
| $Mg^{2+}$ B1. | on hOh B1250 | 2.07Å (0.22Å) |

**Supplementary Figure 4: Active site $Mn^{2+}$ coordination geometry in 2.5Å QPT-1 structure (pdb code: 5cdm) compared with a 2.16Å human top IIβ structure with etoposide (3qx3).**

**(a)** A view of the $Mn^{2+}$ binding active site in 5cdm. **(b)** A final omit map (contoured at 5 sigma) in which oxygens coordinating metal ions have been omittted. Underneath the figure are shown: (i) refined distances (restrained to 2.17Å)and (ii) distances from $Mn^{2+}$ B5081 to peaks in the difference map. Note variability of peak heights and that for two of the waters (B5093 and B5094) there was no prominent difference map peak (indicated by abs. ) Shown is the 2fo-fc 6 sigma (blue) and the fo-fc 9.5 sigma (green). **(c)** A view of the $Mg^{2+}$ binding active site in 3qx3. **(d)** A 'final' omit map (contoured at 7 sigma) in which oxygens coordinating metal ions have been omittted. Underneath the figure are shown: (i) refined distances in 3qx3 (Note est. err is = COORDINATE ERROR (MAXIMUM-LIKELIHOOD BASED) from in 3qx3 from phenix.refine) and (ii) distances from $Mg^{2+}$ B1 to peaks in the difference map. (**e**) the B subunit from re-refined 5cdm is superposed on that in 3qx3; equivalent waters are named. (**f**) the B subunit from 3qx3 is superposed on the re-refined 5cdm structure (e and f are essentially the same).

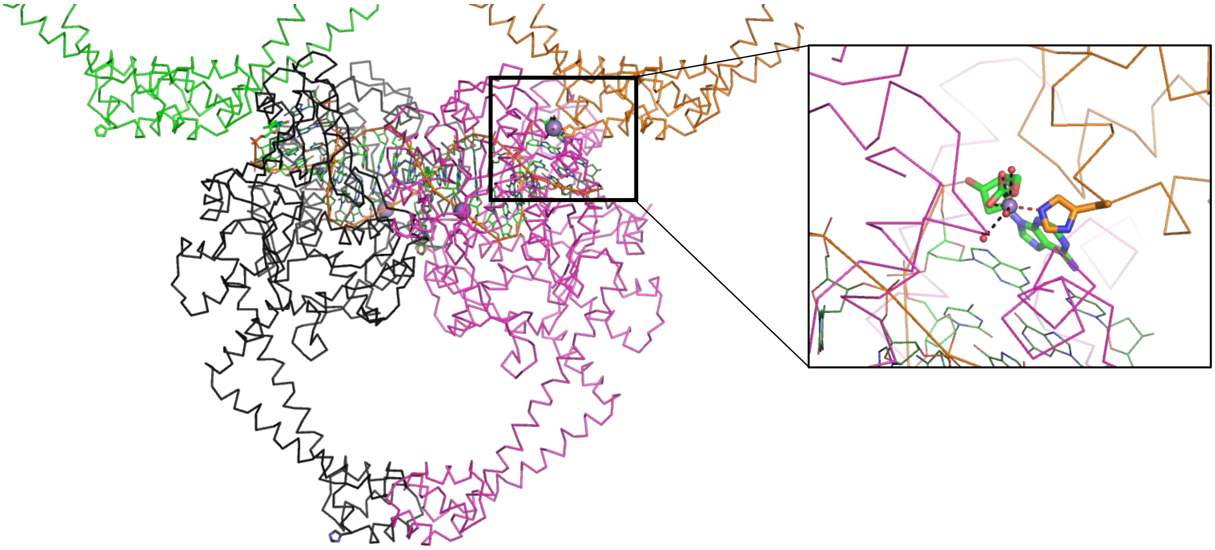

**Supplementary Figure 5: The originally deposited coordinates of 2xcs contained an error at a crystal contact near the Xtal contact $Mn^{2+}$ ion.** In early attempts to understand the crystal packing in the $P6_1$ crystal form an error was accidentally introduced to try and explain observed electron density. The chemistry for the terminal base was wrongly modified. The view is similar to that in Figure 4.
